# Tetherin restricts SARS-CoV-2 replication despite antagonistic effects of Spike and ORF7a

**DOI:** 10.1101/2023.07.28.550997

**Authors:** Elena Hagelauer, Rishikesh Lotke, Dorota Kmiec, Dan Hu, Mirjam Hohner, Sophie Stopper, Mirjam Layer, Rayhane Nchioua, Frank Kirchhoff, Daniel Sauter, Michael Schindler

## Abstract

SARS-CoV-2 infection induces interferon-stimulated genes, one of which encodes Tetherin, a transmembrane protein inhibiting the release of various enveloped viruses from infected cells. Previous studies revealed that SARS-CoV encodes two Tetherin antagonists: the Spike protein (S) inducing lysosomal degradation of Tetherin, and ORF7a altering its glycosylation. SARS-CoV-2 ORF7a has also been shown to antagonize Tetherin. Therefore, we here investigated whether SARS-CoV-2 S is also a Tetherin antagonist and compared the abilities and mechanisms of S and ORF7a in counteracting Tetherin. SARS-CoV and SARS-CoV-2 S reduced Tetherin cell surface levels in a cell type-dependent manner, possibly related to the basal protein levels of Tetherin. In HEK293T cells, under conditions of high exogenous Tetherin expression, SARS-CoV-2 S and ORF7a reduced total Tetherin levels much more efficiently than the respective counterparts derived from SARS-CoV. Nevertheless, ORF7a from both strains was able to alter Tetherin glycosylation. The ability to decrease total protein levels of Tetherin was conserved among S proteins from different SARS-CoV-2 variants (D614G, Cluster 5, α, γ, δ, ο). While SARS-CoV-2 S and ORF7a both colocalized with Tetherin, only ORF7a directly interacted with the restriction factor. Despite the presence of two Tetherin antagonists, however, SARS-CoV-2 replication in Caco-2 cells was further enhanced upon Tetherin knockout. Altogether, our data show that endogenous Tetherin restricts SARS-CoV-2 replication, and that the antiviral activity of Tetherin is partially counteracted by two viral antagonists with differential and complementary modes of action, S and ORF7a.

**IMPORTANCE:** Viruses have adopted multiple strategies to cope with innate antiviral immunity. They blunt signaling and encode proteins that counteract antiviral host factors. One such factor is Tetherin, that tethers nascent virions to the cell membrane and interferes with virus release. For SARS-CoV, the viral glycoprotein Spike (S) and the accessory protein ORF7a are Tetherin antagonists. For pandemic SARS-CoV-2, such activity has only been shown for ORF7a. We therefore analyzed whether SARS-CoV-2 S is a Tetherin-counteracting protein and whether there are differences in the abilities of the viral proteins to antagonize Tetherin. Of note, the efficiency of Tetherin antagonism was more pronounced for S and ORF7a from SARS-CoV-2 compared to their SARS-CoV orthologs. Still, Tetherin was able to restrict SARS-CoV-2 replication. Our results highlight the fundamental importance of the innate immune response in the context of SARS-CoV-2 control and the evolutionary pressure on pathogenic viruses to withhold efficient Tetherin antagonism.

## INTRODUCTION

At the end of 2019, a new coronavirus emerged and spread rapidly around the world, leading the World Health Organization (WHO) to declare it a global viral pandemic on March 11^th^ (1). Its shared sequence identity (79.6%) with severe acute respiratory syndrome coronavirus (SARS-CoV) led to its designation severe acute respiratory syndrome coronavirus 2 (SARS-CoV-2) (2). SARS-CoV-2 belongs to the genus *Betacoronavirus* in the family of *Coronaviridae* and harbors a single-stranded, positive-sense RNA genome (3). Two thirds of the genome encode non-structural proteins (nsps). The remaining 10 kb code for accessory proteins and four structural proteins: nucleocapsid (N), membrane (M), envelope (E) and Spike (S) (4). While N encapsidates the viral genome, E, M and S are incorporated into the host-derived lipid envelope that the virus acquires in the ER-Golgi intermediate compartment (ERGIC), its budding site (5, 6). In contrast, accessory viral proteins such as ORF6 or ORF7a are frequently dispensable for viral replication *in vitro*, but enable efficient replication *in vivo* by suppressing different branches of the host immune response (7).

SARS-CoV-2 infection induces an interferon (IFN) response, which subsequently triggers the expression of interferon-stimulated genes (ISGs) (8, 9). One protein that is induced in most cell types upon exposure to type I IFN is Tetherin (also known as bone marrow stromal antigen 2, BST2, or CD317) (10). It has a unique topology with two membrane anchors, an N-terminal transmembrane domain and a C-terminal glycosylphosphatidylinositol (GPI) anchor (11). These anchors are connected via a glycosylated coiled-coil ectodomain. As a dimer that is linked by disulfide bonds, Tetherin can restrict the release of a variety of enveloped viruses (12). Antiviral activities of Tetherin have been demonstrated for retroviruses (13, 14), rhabdoviruses (15), filoviruses (16), orthomyxoviruses (17), alphaviruses (18), herpesviruses (19), flaviviruses (20) and coronaviruses (21, 22). It is thought that this broad antiviral spectrum is achieved because Tetherin does not recognize specific viral motifs but acts as a physical intermembrane tether. In addition to its eponymous function, Tetherin has been shown to act directly or indirectly as an innate immune sensor for viral infections. It can directly activate NF-κB, thereby inducing a pro-inflammatory signaling response (23, 24). Furthermore, Tetherin-induced endocytosis of tethered viral particles and uptake into endosomes leads to stimulation of endosomal pattern recognition receptors (25).

Many viruses encode Tetherin antagonists with different counteraction strategies. For example, HIV-1 Vpu degrades Tetherin (26, 27), the glycoprotein of Ebola virus has been shown to functionally inhibit Tetherin (28), while SIV Nef proteins and the HIV-2 envelope glycoprotein sequester Tetherin intracellularly in the trans-Golgi network (29–31). For SARS-CoV, two viral proteins have been reported to counteract Tetherin: Spike redirects Tetherin to a lysosomal degradation pathway, leading to decreased Tetherin surface levels (32), and the accessory protein ORF7a binds Tetherin and inhibits its glycosylation, resulting in reduced functionality (33). To date, one report has described SARS-CoV-2 ORF7a as a Tetherin antagonist (22). In addition, SARS-CoV-2 ORF7a was shown to interact with Tetherin (22, 34–36). Whether SARS-CoV-2 Spike (S) is also a Tetherin antagonist and whether there is functional conservation of Tetherin antagonism among S proteins from different viral variants is currently unknown. Furthermore, neither the relative efficiencies of S vs. ORF7a proteins from SARS-CoV and SARS-CoV-2 in antagonizing Tetherin nor potential mechanistic differences have been explored to date. Here, we therefore investigated and directly compared the roles of SARS-CoV-2 S and ORF7a in Tetherin antagonism. We employ functional tests to analyze the effects of the two proteins on surface and total Tetherin levels and determined SARS-CoV-2 replication kinetics in parental versus Tetherin-KO cells. Altogether, our results support a functional role of Tetherin as inhibitor of SARS-CoV-2 despite the presence of at least two viral antagonists.

## METHODS

### Cells and cell culture

Human embryonic kidney (HEK) 293T cells, HEK293T BOI66 MaMTH reporter cells, and cervical cancer-derived HeLa cells were cultivated in DMEM (ThermoFisher) supplemented with 10% fetal calf serum (FCS) and 100 µg/ml Penicillin/Streptomycin (P/S). Human colon cancer-derived epithelial Caco-2 cells were cultured in DMEM containing 10% FCS, 100 µg/ml P/S and 1% non-essential amino acids. Human lung A549 cells were maintained in RPMI medium supplemented with 10% FCS and 100 µg/ml P/S. Caco-2 Tetherin knockdown cell lines were cultured in their cell culture medium with additional 2 µg/ml Blasticidin. All cells were grown in a humidified cell culture incubator at 37°C and 5% CO_2_ and passaged 2 to 3 times a week.

### Generation of Caco-2 Tetherin knockout cell lines

Caco-2 Tetherin knockout cells were generated by lentiviral transduction. Lentiviral stocks were produced in HEK293T cells. To this end, 0.7x 10^6^ HEK293T cells (per well) were seeded in 6-well plates and transfected the next day with 0.45 µg pMD2G, 1.125 µg psPAX2 and 1.5 µg LentiCRISPRv2 blast CD317 (sgRNA sequence: GCTTCAGGACGCGTCTGCAG for KO1, GCTTACCACAGTGTGGTTGC for KO2) using PEI (transfection protocol for HEK293T cells, see below). 24 h post transfection, viruses were harvested. Caco-2 cells were seeded in a 6-well plate at a density of 0.3x 10^6^ cells per well. The supernatant was aspirated one day later and the cells were treated with the lentiviruses produced. 2 days later, the cells were selected with 10 µg/ml blasticidin, and Tetherin KO efficiency was analyzed by Tetherin surface staining and flow cytometry.

### Viruses

Experiments involving replication-competent SARS-CoV-2 were conducted in a BSL3 laboratory. Recombinant SARS-CoV-2 reporter viruses expressing a fluorescent protein instead of ORF6 (SARS-CoV-2 ΔORF6-YFP) (37) or ORF7a (icSARS-CoV-2-mNG) (38) were previously described. IcSARS-CoV-2-mNG was obtained from the World Reference Center for Emerging Viruses and Arboviruses (WRCEVA) at the UTMB (University of Texas Medical Branch). Virus stocks were generated by infection and propagation in Caco-2 cells, and viral titers were determined by infection rates (fluorescent cells) after serial dilution. SARS-CoV-2 ΔORF6-YFP was a kind gift from Prof. Armin Ensser (Institute for Clinical and Molecular Virology, Friedrich-Alexander University Erlangen-Nürnberg (FAU), 91054 Erlangen, Germany).

### Plasmids and antibodies

pEYFP-C1 CD317 was described previously (39). A pmCherry-C1 CD317 construct was created similarly by amplifying and inserting CD317 into pmCherry_C1 via BamHI / XhoI. Plasmids encoding the Spike proteins of SARS-CoV and SARS-CoV-2 were cloned into the pCG IRES-GFP backbone using XbaI / MluI. Human codon-optimized plasmids obtained from Genescript, for SARS-CoV S and from Markus Hoffmann (Infection Biology Unit, German Primate Center – Leibniz Institute for Primate Research, Göttingen, Germany) for SARS-CoV-2 S served as templates. Spike proteins of SARS-CoV-2 variants were generated by amplifying the Spike cDNA sequences from expression plasmids kindly provided by Markus Hoffmann (Infection Biology Unit, German Primate Center – Leibniz Institute for Primate Research, Göttingen, Germany) and inserting them into the pCG-IRES-GFP vector via XbaI / MluI. In all S plasmids, the Kozak sequence (GCCACC) was introduced upstream of the gene. To generate a GFP-Spike fusion protein, SARS-CoV-2 Spike was amplified with primers introducing a 5’-BamHI and 3’-MluI site and allowing an in-frame insertion into the GFP ORF of pWPXLd. Expression plasmids encoding SARS-CoV-1 ORF7a C-V5 and SARS-CoV-2 ORF7a C-V5 were generated by subcloning of inserts into XbaI and MluI restriction sites of the pCG_IRES eGFP vector. SARS-CoV-2 ORF7a was also PCR amplified with primers introducing XhoI / EcoRI restriction sites and cloned into the pmScarlet_C1 backbone. pCG-NL4-3_Vpu-IRES-GFP was described before (40). MaMTH plasmids encoding Gal4, Pex7 and human Tetherin and SARS-CoV-2 Spike were reported before (41, 42). SARS-CoV-2 ORF7a was cloned into the MaMTH Prey vector using Gateway cloning technology (ThermoFisher). The correctness of new constructs was confirmed by Sanger sequencing (Eurofins).

All antibodies used in this study were obtained commercially. Unconjugated antibodies: monoclonal mouse anti-SARS-CoV/SARS-CoV-2 Spike (Biozol, GTX632604), polyclonal rabbit anti-BST2 (Proteintech, 13560-1-AP), polyclonal rabbit anti-V5 (Abcam, ab9116) and monoclonal rat anti-GAPDH (BioLegend, 607902). Conjugated antibodies: PE-conjugated human anti-BST2 (MACS, 130-101-656), goat-anti-mouse 680RD (Li-Cor, 926-68070), goat-anti-rabbit 800CW (Li-Cor, 926-32211), goat-anti-rabbit Alexa594 (Thermo Fisher, A-11012) and goat-anti-rat 800CW (Li-Cor, 926-32219).

### Transfection of HEK293T cells

HEK293T cells were seeded one day prior to transfection in 6-well plates using 7.5×10^5^ cells per well. For transfection of one well, 2 µg plasmid DNA was diluted in 100 µl OptiMEM (ThermoFisher). The DNA mix was incubated with 100 µl OptiMEM containing 6 µl polyethylenimine (PEI) for 15 min at room temperature (RT). The reaction mix was added dropwise to the cells and incubated at 37°C. After 16 h, the medium was changed, and the cells were incubated for another 48 h at 37°C.

### Transfection of HeLa and A549 cells

HeLa and A549 cells were seeded one day prior to transfection in 12-well plates using 2×10^5^ cells per well. For transfection of one well, 100 µl jetPRIME buffer (Polyplus) was mixed with 1 µg DNA and vortexed. After spinning down the DNA mix, 2 µl jetPRIME reagent was added, and the mix was vortexed and spun down again. After 10 min of incubation at RT, the reaction mix was added dropwise to the cells and incubated at 37°C. After 4 h, the medium was changed, and the cells were incubated for another 48 h at 37°C.

### Transfection of Caco-2 cells

Caco-2 cells were seeded one day prior to transfection in 12-well plates using 2×10^5^ cells per well. For transfection of one well, 1 µg plasmid DNA was diluted in 100 µl OptiMEM (ThermoFisher). The DNA mix was incubated with 100 µl OptiMEM containing 3 µl Lipofectamine2000. After 5 min, the reaction mix was added dropwise to the cells and incubated at 37°C. After 4 h, the medium was changed, and the cells were incubated for another 48 h at 37°C.

### Western blot

48 h after the medium was changed following transfection, cells were harvested, washed with PBS and lysed on ice for 30 min using RIPA buffer (10 mM Tris-HCl pH 7.4, 140 mM NaCl, 1% TritonX-100, 0.1% Na-deoxycholate, 0.1% SDS, 1 mM EDTA, 0.5 mM EGTA, Protease Inhibitor Cocktail Tablets, EDTA-Free (Sigma)). Cell lysate was spun down for 10 min at 20.000 x g and 4°C, and the supernatant was mixed with 6x SDS loading buffer and denatured at 95°C for 5 min. Precleared cell lysates were analyzed by sodium dodecyl sulfate polyacrylamide gel electrophoresis (SDS-PAGE) utilizing 10% polyacrylamide gels. Separated proteins were transferred onto a polyvinylidene fluoride (PVDF) membrane by wet transfer. The membrane was blocked in 5% milk in TBS-T (0.1% Tween-20 in TBS) for at least 1 h at RT and subsequently incubated with the primary antibody (in 5% milk in TBS-T) at 4°C overnight or at RT for 2 h. After washing the membrane three times for 10 min with TBS-T, the secondary antibody (in TBS-T) was incubated for 1 h at RT. The membrane was again washed three times with TBS-T, and fluorescence was imaged using the Odyssey® Fc Imaging System (Li-cor).

### Infection experiments

Caco-2 Tetherin knockdown cells and empty-vector transduced control cells were used for infection with SARS-CoV-2. One day prior to infection, 2×10^4^ cells were seeded into a 96-well flat bottom plate. The next day, cells were infected at different multiplicities of infection (MOIs) using infection medium (DMEM, 5% FCS, 100 µg/ml P/S). After 4 h, medium was changed, and viral replication and spread were monitored by live-cell imaging at 37°C for 72 h.

### Fluorescence microscopy

Coverslips were coated with poly-L-lysine (0.01 mg/ml in PBS) for 1 h at 37°C. After washing with PBS, 1.5×10^5^ HEK293T cells were seeded onto the coverslips in a 12-well format. The next day, cells were transfected to express fluorophore-conjugated proteins. After 24 h at 37°C, cells were fixed with 2% PFA in PBS at RT for 15 min. Subsequently, the fixed cells were washed with PBS. To visualize nuclei, cells were stained with DAPI (1:20,000 in PBS) for 10 min at RT. After washing with PBS, the coverslips were transferred onto microscopy slides. The slides were dried overnight at 4°C, and cells were imaged the next day with a fluorescence microscope.

### Immunofluorescence staining and flow cytometry

For flow cytometric analysis, detached cells were resuspended in FACS buffer (1% FCS in PBS) and transferred to a 96-well plate. For cell surface staining, cells were resuspended in PE-conjugated Tetherin antibody dilution (1:11 in PBS) and stained for 30 min at 4°C in the dark. Cells were washed with FACS buffer and stored at 4°C until flow cytometric measurement using the MACSquant VYB cytometer (Miltenyi Biotech). Sinceplasmids encoding GFP via an IRES were used, transfected and un-transfected cells were discriminated via their GFP expression. PE-A was used to gate for PE+, i.e., Tetherin expressing cells. N-fold downregulation was calculated by dividing the mean fluorescence intensity (MFI) of PE in the GFP negative population by the MFI of PE in the GFP positive population. Data were analyzed using the FlowLogic software.

### Mammalian-membrane two-hybrid (MaMTH) assay

HEK293T B0166 *Gaussia* luciferase reporter cells were co-transfected in 96-well plates with 25 ng SARS-CoV-2 protein Bait and 25 ng human Tetherin or control protein Prey MaMTH vectors in triplicates using PEI transfection reagent. Gal4 transcription factor served as a positive control, whereas SARS-CoV-2 Bait proteins with Pex7 Prey were used as negative controls. The following day, Bait protein expression was induced with 0.1 µg/ml doxycycline. Cell-free supernatants were harvested 2 days post-transfection and the released *Gaussia* luciferase was measured 1 s after injecting 20 mM coelenterazine substrate using an Orion microplate luminometer.

## RESULTS

### Coronavirus S and ORF7a downregulate Tetherin in a cell type-dependent manner

To investigate if SARS-CoV- and SARS-CoV-2-derived S and ORF7a proteins modulate Tetherin levels at the plasma membrane, we transfected A549, Caco-2, HeLa and HEK293T cells to express S or ORF7a together with GFP from bicistronic expression plasmids (Fig. 1a). This allowed us to readily quantify Tetherin cell surface levels in the presence of either S or ORF7a via flow cytometry. Of note, the ability of the viral proteins to downregulate Tetherin varied between the four cell lines investigated (Fig. 1a-e). In the lung adenocarcinoma cell line A549 representing human alveolar basal epithelial cells (Fig. 1c), both S proteins downregulated Tetherin about two-fold, whereas in the colon cancer-derived epithelial cell line Caco-2 none of the viral proteins showed a significant effect (Fig. 1d). In contrast, in HeLa cells, only SARS-CoV-2 ORF7a significantly reduced Tetherin cell surface levels (Fig. 1e). To test the effect of basal Tetherin levels on Spike- and ORF7a-mediated downmodulation, we moved to HEK293T cells, which express low endogenous levels of Tetherin (14) and enable an easy titration of Tetherin levels via transfection. Low levels of endogenous Tetherin at the cell surface of HEK293T cells were efficiently downregulated by SARS-CoV and SARS-CoV-2 S (Fig. 1b), similar to the phenotype observed in A549 cells (Fig. 1c). When HEK293T cells were transfected to express increasing amounts of Tetherin, only SARS-CoV-2 S and ORF7a were able to downregulate Tetherin (Fig. 1f), albeit less efficiently than HIV-1 Vpu, an established and highly potent Tetherin antagonist (13).

**Figure 1:**
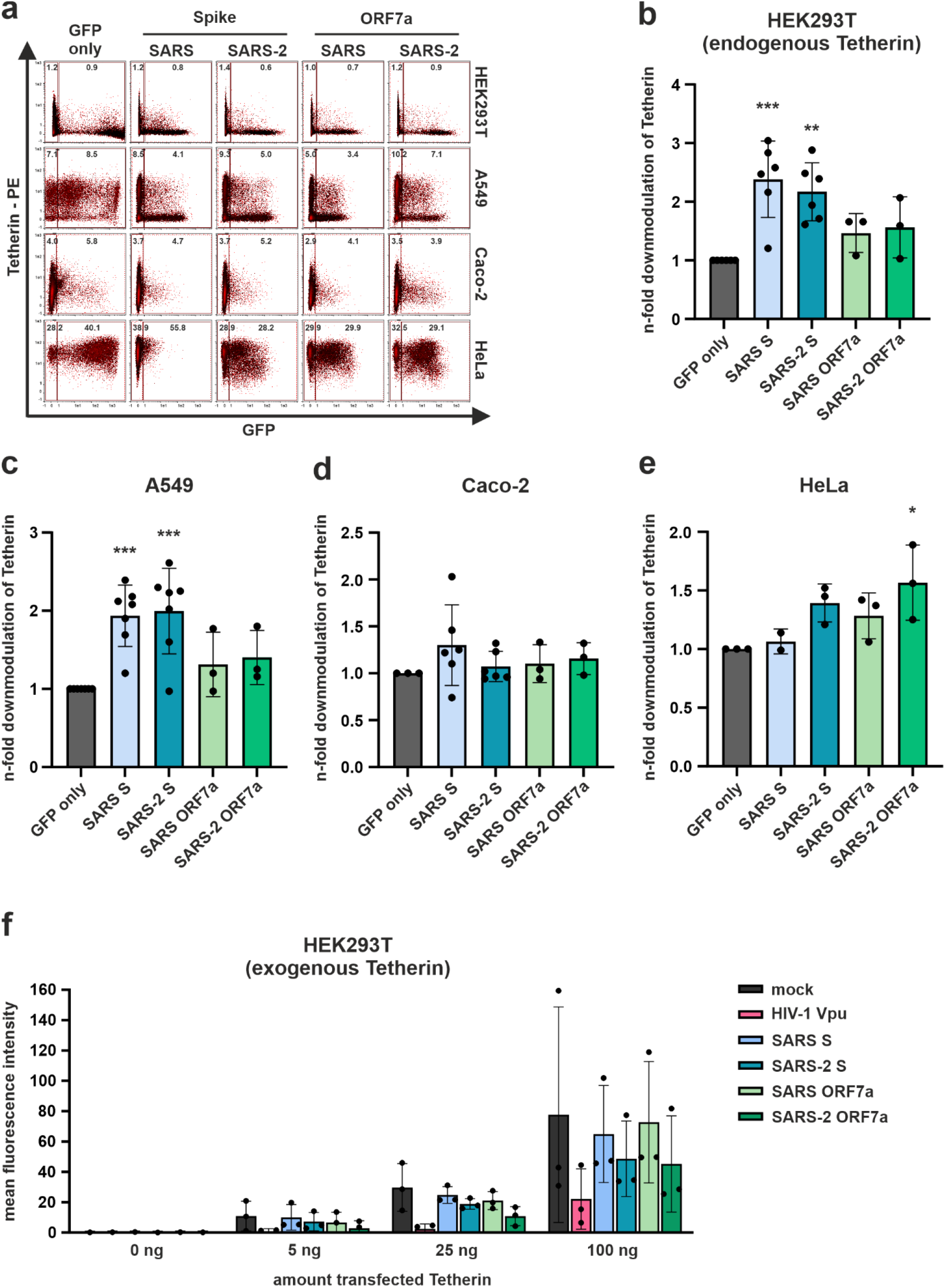
Effect of the SARS-CoV and SARS-CoV-2 Spike (S) or ORF7a protein on Tetherin surface levels in different cell lines. (**a**) HEK293T, A549, Caco-2 and HeLa cells were transfected with expression plasmids encoding for SARS-CoV and SARS-CoV-2 S or ORF7a protein and GFP. Two days post transfection, cells were harvested, surface-stained with a PE-conjugated anti-Tetherin antibody and analysed by flow cytometry. Representative plots of at least three independent experiments are shown and mean PE fluorescence intensity of GFP-negative and GFP-positive population is written. (**b-e**) Mean PE fluorescence of the GFP-negative population was divided by the mean PE fluorescence of the viral protein expressing, GFP-positive population, and the n-fold change compared to the GFP only control was calculated. (**f**) Same experimental setup as in b, but with co-transfection of increasing amounts of Tetherin expression plasmid (5, 25, 100 ng). An expression plasmid for HIV-1 Vpu served as positive control. Mean PE fluorescence is shown. (**b-f**) Mean values of at least three independent experiments are shown. Error bars indicate SD. Statistical significance was tested by one-way ANOVA (*p ≤ 0.05, **p ≤ 0.01, ***p ≤ 0.001).

Together, in the various cell lines tested, either SARS-CoV-2 S or ORF7a downregulated cell surface Tetherin. In contrast, SARS-CoV ORF7a did not show any significant Tetherin-reducing activity, and S from SARS-CoV was only active in A549 and reduced low levels of endogenous Tetherin in HEK293T cells. These data indicate that SARS-CoV-2 evolved at least two antagonists that differentially act on Tetherin, dependent on the cellular context and the overall protein levels of this restriction factor.

### SARS-CoV-2 S and ORF7a reduce total cellular Tetherin levels

Thus far, we analyzed the ability of the viral proteins to reduce levels of cell surface Tetherin, assuming that expression of Tetherin at the plasma membrane is important for its antiviral activity. However, SARS-CoV and SARS-CoV-2 bud into intracellular vesicles (5, 6) and may therefore be trapped and restricted by Tetherin intracellularly. Furthermore, there could be differential effects of viral antagonists on *de novo* synthesized versus cell surface Tetherin. We hence used the established HEK293T Tetherin expression model (Fig. 1f), but instead of measuring Tetherin at the cell surface by flow cytometry, cellular lysates were prepared, and total Tetherin levels were analyzed and quantified by Western blot (WB) (Fig. 2). Using this approach, we found that SARS-CoV-2 S and ORF7a, and to a lesser extent SARS-CoV S, reduced total cell-associated Tetherin levels (Fig. 2a and b). Of note, the ORF7a proteins of both viruses also affected Tetherin glycosylation, as inferred from a strongly altered Tetherin migration pattern under denaturating conditions in the presence of ORF7a (Fig. 2a). Similarly, we tested the ability of SARS-CoV-2 S variants (α, γ, δ, ο, D614G, Cluster 5) to antagonize Tetherin. This revealed that S proteins derived from all tested variants retained the ability to reduce total Tetherin levels (Fig. 2c and d), indicating that there is an ongoing selection pressure on this S protein activity.

**Figure 2:**
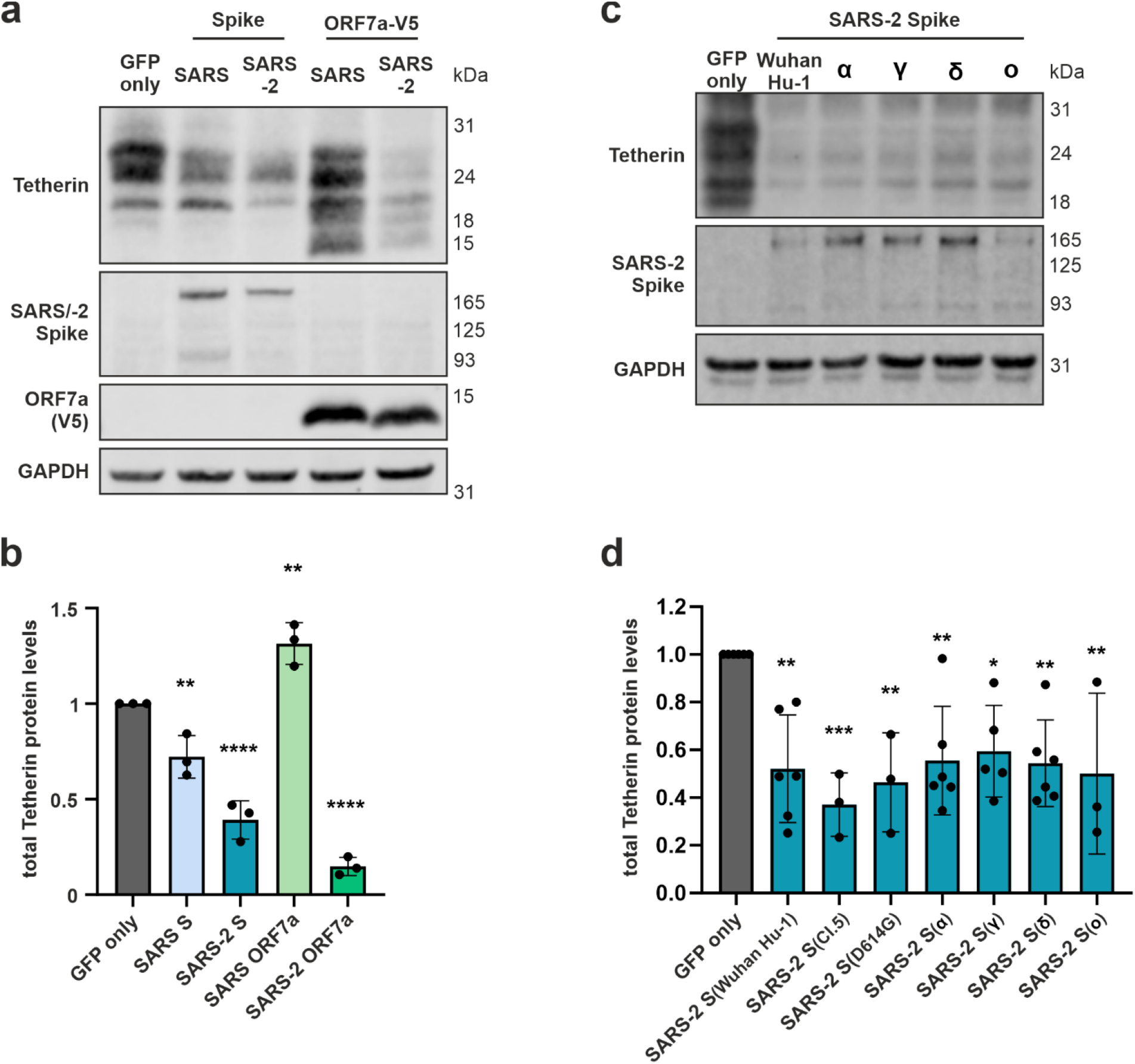
Effect of the SARS-CoV or SARS-CoV-2 Spike (S) and ORF7a protein on total Tetherin levels. (**a**) HEK293T cells were co-transfected with expression plasmids encoding for Tetherin and the SARS-CoV or SARS-CoV-2 S or ORF7a protein. Two days post transfection (d.p.t.), cells were harvested and lysed. Total Tetherin protein levels in cell lysates were determined by Western blot. (**b**) Tetherin levels were quantified and normalized to GAPDH. Tetherin levels of the GFP only control were set to 1. (**c**) HEK293T cells were co-transfected with expression plasmids encoding for Tetherin and S proteins of different SARS-CoV-2 variants. Two d.p.t., cells were harvested, and cell lysates were analyzed by Western blot. (**d**) Tetherin levels were quantified and normalized as described in b. (**a, c**) Representative blot of three independent experiments. (**b, d**) Mean values of at least three independent experiments. Error bars indicate SD. Statistical significance was tested by one-way ANOVA (*p ≤ 0.05, **p ≤ 0.01, ***p ≤ 0.001).

### SARS-CoV-2 S and ORF7a colocalize with Tetherin, but only ORF7a shows evidence for interaction with the restriction factor

We next sought to assess a potential interaction between Tetherin and SARS-CoV-2 S or ORF7a. To this end, we transfected HEK293T cells to co-express Tetherin fused to mCherry or YFP, as well as either SARS-CoV-2 S fused to GFP or ORF7a fused to mScarlet. Then, cells were analyzed for protein localization by confocal fluorescence microscopy. Upon co-expression of Tetherin and the two SARS-CoV-2 proteins, we detected pronounced areas of co-localization, indicating that S and ORF7a are in close proximity to Tetherin (Fig. 3). To further elucidate if the viral proteins directly interact with Tetherin, we took advantage of the mammalian-membrane two-hybrid (MaMTH) assay (Fig. 4). In this assay, the transcription factor Gal4 is released from a bait protein upon interaction with its prey, due to reconstitution of ubiquitin that is cleaved by deubiquitinating enzymes. Then, Gal4 activates luciferase expression in the nucleus (Fig. 4a). Over-expression of Gal4 served as positive control, while a random protein (Pex7) served as negative control. As expected, expression of individual prey (Tetherin) or bait (S, ORF7a) constructs resulted in low luciferase reporter activity as compared to Gal4 only expression (Fig. 4b). Similarly, luciferase signals were close to the Pex7 negative control when SARS-CoV-2 S bait was combined with Tetherin prey (Fig. 4c). Importantly, however, the combination of ORF7a with Tetherin resulted in a robust luciferase signal that was comparable to the positive control Gal4 (Fig. 4c). We hence conclude that, while both SARS-CoV-2 S and ORF7a share areas of subcellular co-localization with Tetherin, based in our assay system, only ORF7a shows evidence for a physical interaction with the antiviral factor.

**Figure 3:**
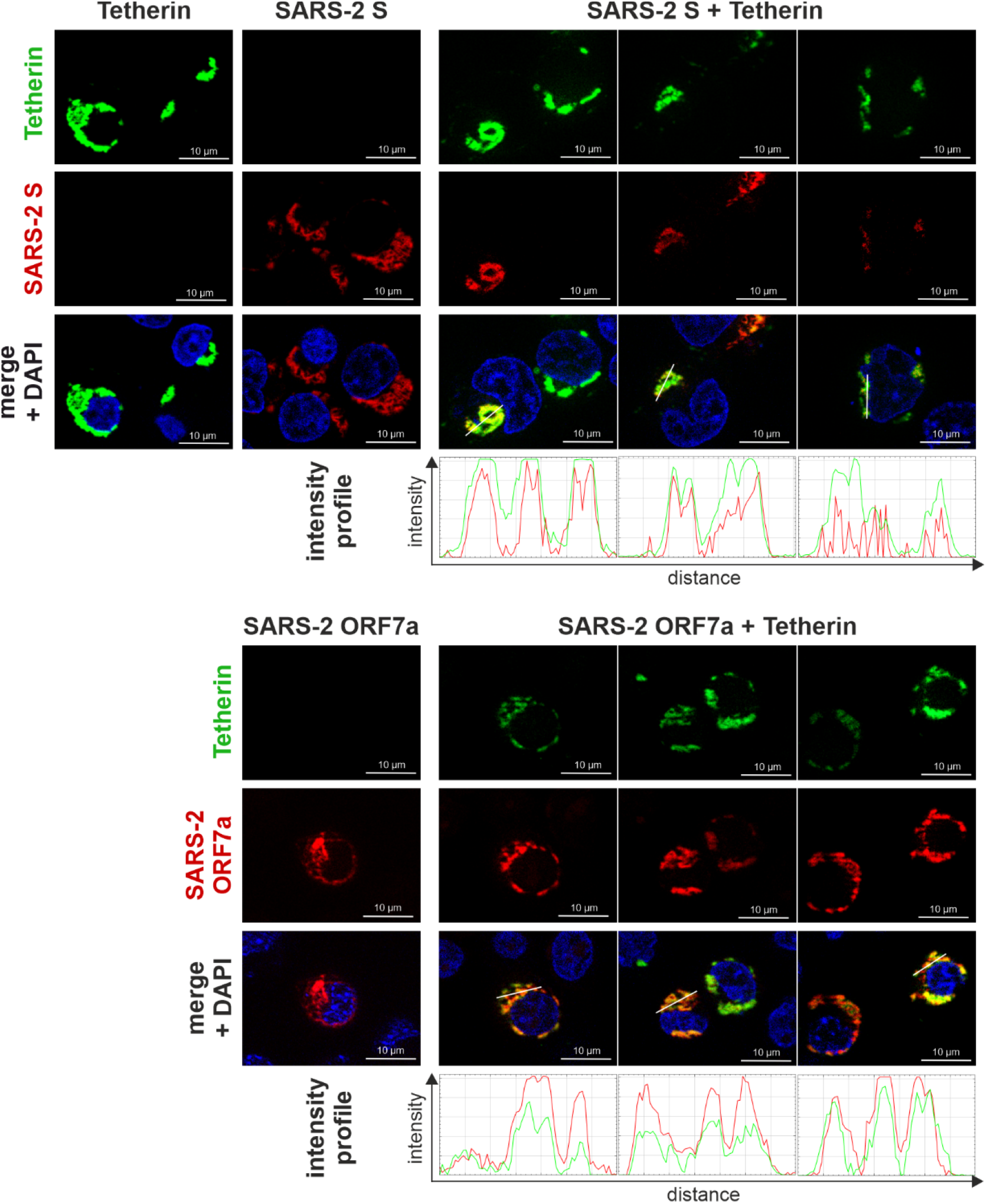
SARS-CoV-2 Spike (S) and ORF7a colocalize with Tetherin. HEK293T cells were transfected with expression plasmids encoding SARS-CoV-2 S-GFP or SARS-CoV-2 ORF7a-mScarlet and Tetherin-mCherry or Tetherin-YFP. For simplicity, Tetherin is shown in green and the viral proteins in red. 24 hours post transfection, cells were fixed with 2% PFA and stained with DAPI. Images were acquired using a fluorescent microscope and processed using ImageJ. For colocalization analysis, intensity profiles for indicated lines are shown. Representative images of two independent experiments are shown (scale bar = 10 µm).

**Figure 4:**
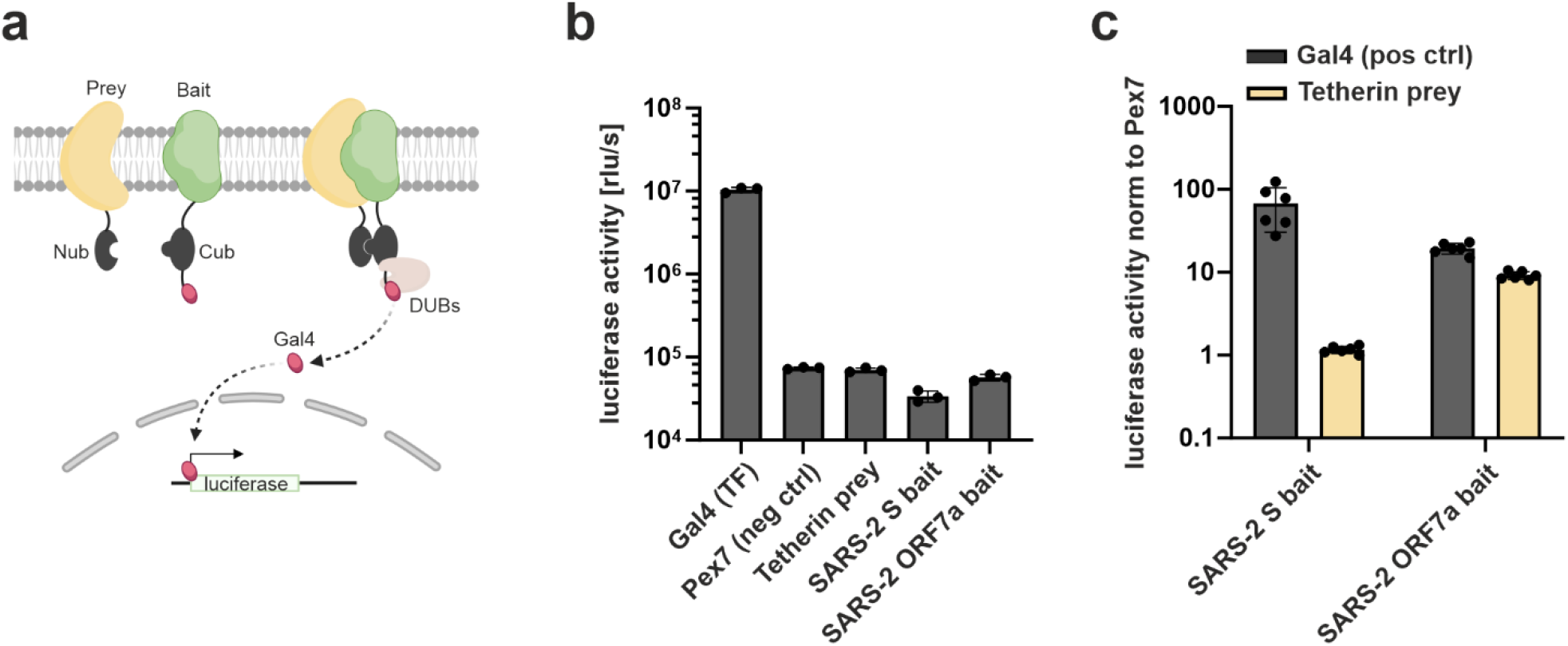
Mammalian-membrane two-hybrid assay (MaMTH) to analyze Tetherin interactions with SARS-2 Spike (S) or ORF7a. (**a**) Schematic representation of the principle of the MaMTH assay. Bait and prey proteins are tagged with two inactive fragments of ubiquitin. Interaction of the bait and prey protein leads to reconstitution of an active ubiquitin that is cleaved by endogenous deubiquitinating enzymes (DUBs). This leads to the release of the transcription factor Gal4 that activates reporter gene (*Gaussia* luciferase) transcription. The scheme was created using BioRender, adapted from Saraon et al. (51). (**b, c**) BOI66 HEK293T reporter cells are transfected with prey and bait constructs. Expression plasmids for Gal4 and Pex7 served as positive and negative controls, respectively. After 48 h, cell culture supernatants were harvested and luminescence was measured. (**b**) Luciferase activity of controls is shown. (**c**) Tetherin was used as prey combined with SARS-2 S or ORF7a as bait. The transcription factor GAL4 was again added as a positive control. Luciferase activity was normalized to the Pex7 negative control. Mean values of at least three independent experiments are shown. Error bars indicate SD.

### Tetherin restricts replication of authentic SARS-CoV-2

We next aimed to assess the importance for SARS-CoV-2 to maintain functional Tetherin antagonism in the context of viral infection of SARS-CoV-2 permissive cells. For this, we decided to monitor viral replication in the presence and absence of Tetherin. These experiments were conducted in Caco-2 epithelial cells, since those cells sustain authentic SARS-CoV-2 replication without the necessity of overexpressing the entry receptor ACE2. Furthermore, Caco-2 cells express considerable levels of intracellular endogenous Tetherin (43) that might hence localize to the internal compartments of SARS-CoV-2 assembly and release. Upon CRISPR/Cas9-mediated knock-out of Tetherin in these cells (Fig. 5a and b), we infected two independent bulk transduced heterogenous cell lines with SARS-CoV-2 reporter viruses either expressing yellow fluorescent protein (YFP) instead of ORF6 or mNeonGreen (mNG) instead of ORF7. We monitored viral replication and spread by live cell imaging for 72 h (Fig. 5c-f). Of note, in both Tetherin KO cell lines we observed increased spread and replication of both reporter viruses compared to control cells, especially at ∼24 h to ∼48 h at different MOIs (Fig. 5c and d). This difference turned out to be significant for the SARS-CoV-2 ΔORF6-YFP infected cells, when total fluorescence was integrated as area under the curve (AUC) as a proxy for overall viral replication (Fig. 5e) and followed the trend for the SARS-CoV-2 ΔORF7-mNG reporter virus (Fig. 5f). Altogether, endogenous Tetherin restricts replication of SARS-CoV-2 in epithelial cells, in the presence of at least two (ΔORF6) or one (ΔORF7) functional Tetherin antagonists.

**Figure 5:**
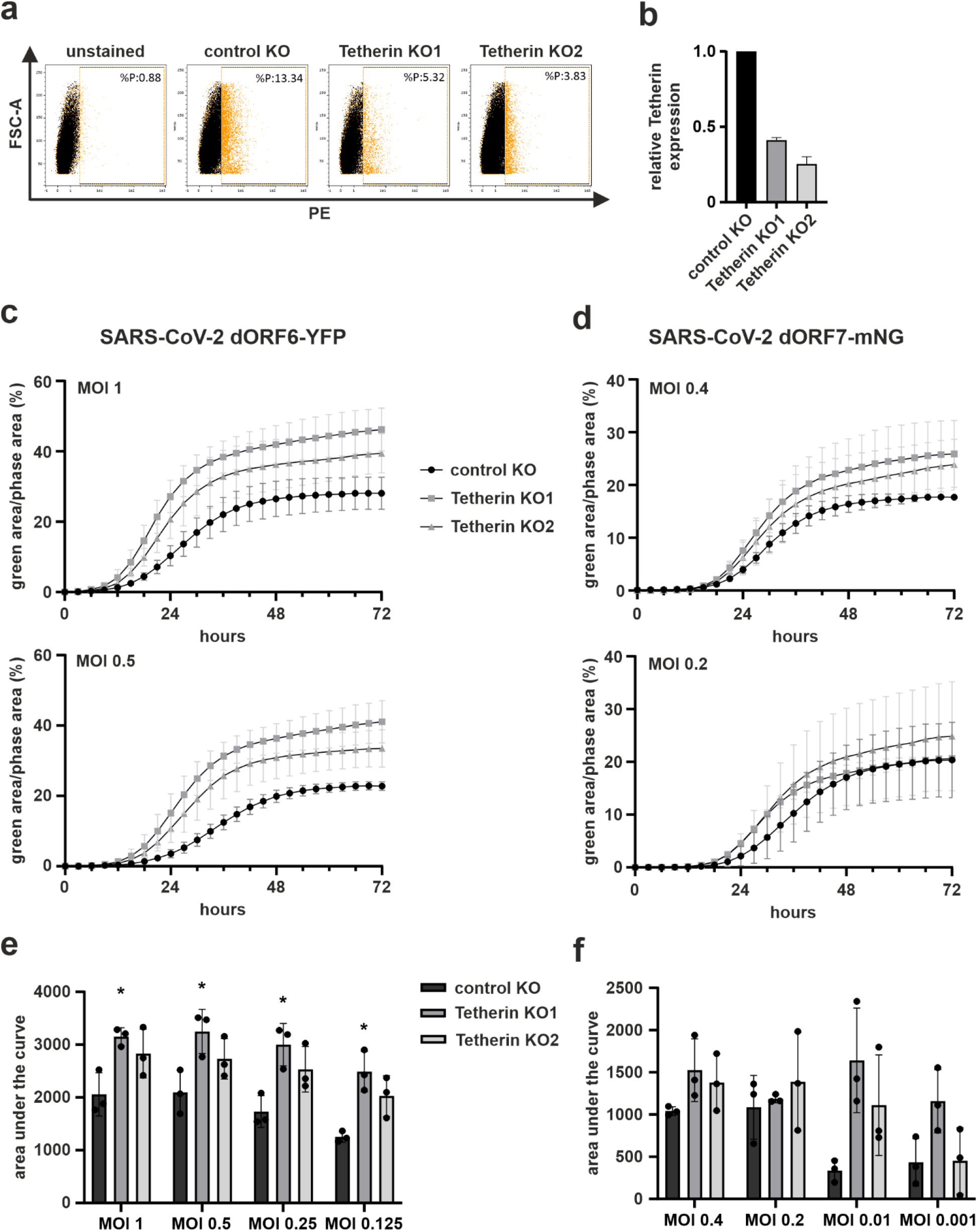
Orf6- and orf7-deleted SARS-CoV-2 replication is increased in Tetherin knockdown Caco-2 cells. (**a**) Validation of Tetherin knockdown. Caco-2 control and knockdown cells (KO1, KO2) were stained by cell surface staining using a PE-conjugated anti-Tetherin antibody. Tetherin expressing cells are shown in orange. (**b**) Percentage of Tetherin positive (PE+) cells normalized to control cells. **(c, d)** Caco-2 control and Tetherin knockdown cells were infected with (**c**) SARS-CoV-2 ΔORF6-YFP or (**d**) SARS-CoV-2 ΔORF7-mNG at different MOIs. Viral replication rates and spread, evident by fluorescent signal, were monitored for 72 h by live-cell imaging. Quantification of the areas under the curve for (**e**) SARS-CoV-2 ΔORF6-YFP or (**f**) SARS-CoV-2 ΔORF7-mNG. (**c-f**) Mean values of three independent experiments conducted in triplicates are shown. Statistical significance was tested by one-way ANOVA (*p ≤ 0.05).

## Discussion

Coronaviruses including SARS-CoV-2 induce an interferon response that is counteracted at several levels (44, 45). One viral strategy is to encode antagonists of specific ISGs and this has been demonstrated for Tetherin, which otherwise restricts release of assembled virus particles, for instance HIV-1 (13, 14). Examples of known viral Tetherin antagonists are the lentiviral accessory proteins Vpu and Nef (26, 27, 46–48), KSHV K5 (49, 50) or the Ebola virus glycoprotein (28). For SARS-CoV and SARS-CoV-2, accumulating evidence suggests that ORF7a is a Tetherin antagonist that can directly bind and alter the glycosylation pattern of Tetherin (22, 33). Similarly, one report suggested that the SARS-CoV S protein acts as Tetherin antagonist (32). Here, we confirm this observation and extend it to the S protein of SARS-CoV-2. While SARS-CoV and SARS-CoV-2 S proteins actively reduce cell surface levels of Tetherin in various cell lines, no or only marginal such activity was observed for ORF7a, which is in line with differential and thus complementary modes of Tetherin antagonism by the two viral proteins (Fig. 1). Furthermore, both S and ORF7a modulated total Tetherin levels, albeit to different extents. While S and ORF7a of SARS-CoV only marginally reduced total cellular Tetherin, SARS-CoV-2 S potently reduced total Tetherin protein levels, an activity that was also inherent to SARS-CoV-2 ORF7a (Fig. 2a and b). In addition, ORF7a derived from SARS-CoV and SARS-CoV-2 altered Tetherin glycosylation (Fig. 2a). In conclusion, both, SARS-CoV and SARS-CoV-2 evolved Tetherin antagonism by at least two viral proteins. However, SARS-CoV-2 S is able to efficiently reduce both total and surface Tetherin, and ORF7a from SARS-CoV-2 alters glycosylation as well as total levels of cellular Tetherin. In contrast, S and ORF7a from SARS-CoV are only able to reduce cell surface Tetherin and modulate its glycosylation, respectively. At present, our data do not show major differences in the ability of SARS-CoV-2 variant-specific S proteins to reduce total cellular Tetherin levels (Figure 2b and d). Hence, Tetherin antagonism seems to be a conserved function of SARS-CoV-2 S. Whether ORF7a proteins derived from SARS-CoV-2 variants differ in their ability to counteract Tetherin is currently unknown and subject to further investigation.

Another important finding of our work is the notion that Tetherin is still able to restrict SARS-CoV-2 replication in the presence of the two Tetherin antagonists S and ORF7a, as shown by our live cell replication kinetics with SARS-CoV-2-ΔORF6-YFP (Fig 5 c and e) in Caco-2 cells with and without Tetherin expression. This highlights the need for SARS-CoV-2 to withhold additional mechanisms that blunt the interferon response, not only at the level of effector proteins, but also during induction. Furthermore, SARS-CoV-2 lacking ORF7a has similar replication kinetics as compared to the variant expressing both Tetherin antagonists. In conclusion, at least in our model, S can compensate for ORF7a to antagonize Tetherin, indicating a high pressure on viral evolution to maintain functional Tetherin antagonism.

In conclusion, we here confirm and extend previous findings on Tetherin counteraction by SARS-CoV and the pandemic SARS-CoV-2. We demonstrate functional Tetherin antagonism by the S and ORF7a proteins of both viruses. Moreover, there are differences in the mechanisms and efficencies of Tetherin antagonism, when comparing S and ORF7a of SARS-CoV and SARS-CoV-2. Whether the latter contribute to the higher transmissibility, spread and pathogenicity of SARS-CoV-2 is an important question that warrants further investigation.

### Consent for publication

All authors gave their consent to publish. All authors read and approved the final manuscript.

## Availability of data and material

All data generated and analyzed during this study are included in this published manuscript.

## Competing Interests

The authors declare that they have no competing interests.

## Funding

DS and MS were supported by a COVID-19 research grant of the Federal Ministry of Education and Research (MWK) Baden-Württemberg. DS was supported by the Heisenberg Program of the German Research Foundation [grant number SA 2676/3-1]. FK is supported by the DFG (CRC 1279) and an ERC Advanced grant (Traitor-viruses). DK and RN are supported by a Baustein grant from Ulm University.

## Authors’ contributions

EH performed most of the experiments, including flow cytometric measurements of Tetherin modulation and western blot analyses of Tetherin expression, colocalization analyses and importance of endogenous Tetherin for SARS-CoV-2 replication. DK and RN performed the MaMTH assay. RL, DH and ML supported experiments done by EH, and FK studies performed by RN and DK. MH, SS and DH generated Tetherin and SARS viral protein expression plasmids. Tetherin knockout cell lines were generated by ML, supported by DH. EH and MS planned and designed most of the experiments and analyzed the data, supported by RL and DS. DS, FK and MS provided reagents and analysis tools. EH and MS wrote the initial manuscript draft. All authors contributed to editing and developed the manuscript to its final form.

## Acknowledgements

We thank Markus Hoffmann for providing expression plasmids (Leibniz Institute for Primate Research, Göttingen, Germany) and Armin Ensser (Virology, University Hospital Erlangen) for providing the SARS-CoV-2 ΔORF6-YFP.

